# Common allotypes of ER aminopeptidase 1 have substrate-dependent and highly variable enzymatic properties

**DOI:** 10.1101/2020.11.19.389403

**Authors:** Jonathan P. Hutchinson, Ioannis Temponeras, Jonas Kuiper, Adrian Cortes, Justyna Korczynska, Semra Kitchen, Efstratios Stratikos

## Abstract

**Objective:** Polymorphic variation of immune system proteins can drive variability of individual immune responses. ER aminopeptidase 1 (ERAP1) generates antigenic peptides for presentation by MHC class I molecules. Coding single nucleotide polymorphisms (SNPs) in *ERAP1* have been associated with predisposition to inflammatory rheumatic disease and shown to affect functional properties of the enzyme, but the interplay between combinations of these SNPs as they exist in allotypes, has not been thoroughly explored.

**Methods:** We used phased genotype data to estimate ERAP1 allotype frequency in 2,504 individuals across five major human populations, generated highly pure recombinant enzymes corresponding to the 10 most common ERAP1 allotypes and systematically characterized their *in vitro* enzymatic properties.

**Results:** We find that ERAP1 allotypes possess a wide range of enzymatic activities, up to 60-fold, whose ranking is substrate-dependent. Strikingly, allotype 10, previously associated with Behçet’s disease, is consistently a low-activity outlier, suggesting that a significant percentage of individuals carry a sub-active ERAP1 gene. Enzymatic analysis revealed that ERAP1 allotypes can differ in both catalytic efficiency and substrate affinity, differences that can change intermediate accumulation in multi-step trimming reactions. Alterations in efficacy of an allosteric inhibitor that targets the regulatory site suggest that allotypic variation influences the communication between the regulatory and the active site.

**Conclusion:** Our work defines the wide landscape of ERAP1 activity in human populations and demonstrates how common allotypes can induce substrate-dependent variability in antigen processing, thus contributing, in synergy with MHC haplotypes, to immune response variability and to predisposition to chronic inflammatory conditions.

## Introduction

Major Histocompatibility Complex molecules (MHC, Human Leukocyte Antigens in humans) are the most polymorphic human genes with tens of thousands different allomorphs identified to date [1]. MHC class I molecules (MHCI), bind small protein fragments (peptides) that originate from normal cellular proteins or pathogen proteins and then translocate to the cell surface to present their cargo to cytotoxic T-lymphocytes [2]. Polymorphic variation in MHCI predominantly affects the structure of the binding groove and allows the presentation of a large variety of peptide sequences.

MHCI bind their peptide cargo in the Endoplasmic Reticulum (ER) with the assistance of the peptide loading complex [3]. While MHCI tend to bind peptides that are between 8-11 amino acids long (the majority of which are 9mers) many peptides that enter the ER can be substantially longer [4]. Two ER-resident aminopeptidases, ER aminopeptidase 1 and ER aminopeptidase 2 (ERAP1 and ERAP2) catalytically process these precursor peptides and define the peptide pool that is available for binding onto MHCI [5].

The *ERAP1* gene is also polymorphic and a variety of coding single nucleotide polymorphisms (SNPs) confer susceptibility to human disease, most notably chronic inflammatory conditions, often in epistasis with HLA class I alleles, which emphasizes the critical role of ERAP1 in antigen presentation [6-9]. The genetic association inflammatory diseases such as *HLA-B27-associated Ankylosing Spondylitis, HLA-B51-associated Behçet’s Disease* and *HLA-A29-associated Birdshot Uveitis* led to the hypothesis that these conditions are driven by pathogenic changes in antigen presentation as a direct result of alterations in substrate preferences or activity of ERAP1 [10-13]. Several ERAP1 SNPs have been described to affect the function of the enzyme [14, 15]. Mechanisms proposed to underlie this effect include direct interactions with the substrate [16], effects on conformational dynamics [17], protein expression level [18, 19], or combinations of these[10]. However, not all of the possible combinations of the 9 most common coding SNPs (i.e. allotypes) [20] occur at equal frequency in the human population [9]. Rather, these SNPs encode a limited palette of allotypes that are maintained at high frequency (>1%) in populations, which suggests functional asymmetry between ERAP1 allotypes. This is supported by the fact that some ERAP1 allotypes are protective while others confer risk to inflammatory diseases [21]. A deep understanding of the functional properties of ERAP1 allotypes rather than individual SNPs is critical to unraveling their physiological impact on disease.

Previous studies have described several ERAP1 allotypes. Most studies, defined ERAP1 allotypes as the combination of 9 coding SNPs at aminoacid positions 56, 127, 276, 346, 349, 528, 575, 725 and 730. Ombrello et al reported ten common ERAP1 allotypes in three populations of European and East Asian Ancestry (n=160) [9]. Reeves et al reported some additional distinct allotypes discovered in small patient cohorts of Ankylosing Spondylitis (n=72) [15] and oropharyngeal squamous cell carcinoma (n=25) [22]. These allotypes however have been controversial and proposed to be rare by others [23, 24]. While these studies have contributed to our understanding of the emerging role of ERAP1 allotypes in disease, controversy still remains on which ERAP1 allotypes are common and systematic analysis of their functional differences has been lacking.

We used phased genotype data from the 1000 genome project [25] to define common ERAP1 allotypes in 2504 individuals of five major human populations. We generated recombinant versions of the 10 most common allotypes and comprehensively characterized their *in vitro* enzymatic properties. We find a complex landscape of large substrate-dependent enzymatic activity differences between allotypes due to effects on catalytic efficiency and substrate affinity. Our findings suggest that ERAP1 allotypic variation has the potential to strongly synergize with MHCI alleles, in an epitope dependent manner, to enhance immune system variability in natural human populations.

## Experimental methods

### Materials

Leu-pNA and Leu-AMC were purchased from Sigma-Aldrich. Peptide with the sequence LLKHHAFSFK was purchased from Genecust. Peptide LLRIQRGPGRAFVTI was purchased from JPT peptide technologies. Peptide YTAFTIPSI was purchased from BioPeptide Inc. and ovalbumin peptides were from CRB Discovery. All peptides were HPLC-purified to >95% purity and confirmed by mass spectrometry to have the correct mass. Inhibitor DG013A was synthesized as described previously [26] and inhibitor GSK849 was obtained, purified and characterized as described previously [16].

### ERAP1 allotype estimation in the human population

Available phased genotype data for 9 coding SNPs at 5q15 in 2504 samples of 26 ethnic groups of European (EUR), African (AFR), East Asian (EAS), South Asian (SAS), and mixed American (AMR) ancestry were obtained from the 1000 Genomes Project Phase 3 [25]. Phased genotypes were used to estimate the allotypes on each chromosome based on 9-SNP haplotypes, which occur in >1 % of all populations. Combinations of the estimated allotype frequencies were plotted using the R package *ggplot2 [27]*.

### Gene constructs, protein expression and purifications

The sequence corresponding to full length ERAP 1 allotype 2 with a non-cleavable C-terminal 6 His-tag was codon optimised for insect expression and the gene was chemically synthesised. The gene was then ligated with the BamHI and XhoI-linearised pFASTBAC1 vector using T4 ligase. The product of the ligation reaction was transformed into the competent cells and the positive clones were selected by single colony screening and DNA sequencing. The positive plasmid was further verified by digestion with BamHI and XhoI. The remaining ERAP1 allotypes were generated by site directed mutagenesis using allotype 2 construct as a DNA template. Protein expression was performed as previously described, with the exception that the C-terminal His tag was not removed [28]. Concentrations of protein stock solutions were determined spectrophotometrically using an extinction coefficient of 171,200 M^−1^cm^−1^ at 280 nm.

### Enzymatic assays with dipeptide substrates

The enzymatic activity of ERAP1 was measured using the dipeptidic fluorigenic substrate Leucine-7-amido-4-methycoumarin (Leu-AMC) as previously described [29]. Briefly, the change in fluorescence at 460 nm (excitation at 380 nm) was followed over time using a TECAN SPARK 10m plate reader. A standard curve of AMC was used to convert the signal to product concentration. For MM measurements the dipeptide substrate L-Leucine-para-nitroanilide (Leu-pNA) was used and the generation of pNA was followed by measuring the absorption at 405 nm as described previously [29]. For experiments measuring the rate of hydrolysis versus enzyme or substrate concentrations, measurements using the dipeptidic fluorogenic substrate Leucine-7-amidocoumarin (Leu-AMC) were made in a buffer of 20 mM HEPES pH 7.0, 100 mM NaCl, 0.002% Tween 20, in deep well 384 well plates at a final volume of 50 uL/well at 20°C, using a Tecan M1000 plate reader with 360 nm excitation and 460 nm emission (5 nm bandpass on both monochromators). Measurements where enzyme concentration was varied were set up using a multichannel pipette and initiated by adding a final concentration of 25 μM Leu-AMC. Measurements where substrate concentration was varied were set up using a Hewlett Packard D300 digital dispenser and initiated by adding a final concentration of 25 nM ERAP1 (allotypes 1-9) or 250 nM ERAP1 (Allotype 10). An aminocoumarin (AMC) standard curve was used to convert fluorescence intensity to product concentration, after which initial rates were obtained by linear fit of the early region of the timecourses.

### Enzymatic assays with peptides

Trimming of peptides LLRIQRGPGRAFVTI and LLKHHAFSFK was performed as described previously [16]. Briefly, 20 μM peptide and 1 nM enzyme at a final assay volume of 200 μL were mixed in assay buffer of 20 mM Heres pH 7.0, 100 mM NaCl, 0.002% Tween 20 and incubated for 30 min at 37 °C. Reactions were carried out in 3 replicates for each allotype, stopped by freezing and were stored at −80 °C until analyzed by HPLC chromatography. For trimming assays using the YTAFTIPSI substrate, measurements were made in an assay buffer of 20 mM Hepes pH 7.0, 100 mM NaCl, 0.002% Tween 20, in deep well 384 well plates at a final assay volume of 25 uL/well and temperature of 20°C. For the Michaelis-Menten kinetics a range of concentrations of YTAFTIPSI peptide was dispensed using a Hewlett Packard D300 digital dispenser and reactions were initiated by addition of a final concentration of 1 nM ERAP1 (allotypes 1-9) or 10 nM ERAP1 (allotype 10). Reactions were stopped after 60 minutes incubation by addition of 25 μL of 0.75 % TFA in water containing 5 μM Ac-YTAFTIPSI as internal standard. Mass signal intensities corresponding to the product (TAFTIPSI) and internal standard (Ac-YTAFTIPSI) were measured on a Rapidfire autosampler (Agilent) equipped with a C18 solid phase cartridge, coupled to a Sciex 4000 Q-trap MS (AB Sciex), using Multiple reaction monitoring (MRM), as described previously [30]. Integrated product intensity signals were normalized to the respective internal standard intensity signals, before conversion to product concentration using a TAFTIPSI standard curve. Turnover did not exceed 35% at any substrate concentration. Data were fitted to the Michaelis-Menten equation as described previously [30]. To measure inhibition by GSK849, a 3-fold dilution series of inhibitor was made in DMSO and 250 nL of each concentration point was dispensed to a 384 well assay plate using an Echo acoustic dispenser. ERAP1 was added followed by YTAFTIPSI substrate at a final concentration of 5 uM. Final ERAP1 concentrations varied according to allotype as follows: 0.5 nM allotype 3; 1 nM allotypes 1,2,4,5,6,8 and 9; 2 nM allotype 7; 10 nM allotype 10. After incubation for 60 minutes, reactions were stopped and measured as described above. Calibration curves were measured for substrate (YTAFTIPSI) and product (TAFTIPSI), and used to correct for the difference in detection sensitivity between the two analytes. The corrected intensity data were used to calculate % turnover, from which product concentration was calculated. Data were normalized to % enzymatic activity between high (uninhibited) and low (100 μM DG13) controls and fitted to a 4 parameter logistic expression as described previously [30].

To follow the trimming of ovalbumin epitope precursors, measurements were made in an assay buffer of 50 mM Hepes pH 7.0, 100 mM NaCl, 0.002% Tween 20, in deep well 384 well plates at a final assay volume of 50 μL/well and temeprature of 20°C. ERAP1 (allotypes 1-9 at 10 nM final concentration, allotype 10 at 100 nM final concentration) were mixed with 14 mer peptide GLEQLESIINFEKL (at 25 μM final concentration). Wells were stopped at increasing time points by the addition of 50 μL of 0.75% TFA in water containing 5 μM Ac-YTAFTIPSI as internal standard. Mass signal intensities corresponding to the 7 ovalbumin peptide species (14-mer GLEQLESIINFEKL, 13-mer LEQLESIINFEKL, 12-mer EQLESIINFEKL, 11-mer QLESIINFEKL, 10-mer LESIINFEKL, 9-mer ESIINFEKL, 8-mer SIINFEKL) and internal standard (Ac-YTAFTIPSI) were measured using the Rapidfire device as described above. Signals at each peptide mass were divided by the respective internal standard signal. Normalized data were converted to peptide concentration using calibration curves for each peptide species.

## Results

Analysis of the Genome Aggregation Database that contains 125,748 exome sequences [31] using a 1% frequency cutoff to qualify a coding genomic missense variant as a polymorphism, revealed only 10 amino acid positions as polymorphic, namely 12, 56, 127, 276, 346, 349, 528, 575, 725 and 730, consistent with a previous study [9]. Since however position 12 lies in the signal peptide that is normally excised after translocation of ERAP1 into the ER and thus does not appear in the mature protein, we focused our analysis on the remaining 9 positions. These 9 SNPs could be theoretically organized in up to 2^9^ discreet allotypes. To define which ERAP1 allotypes are common in human populations we exploited available phased (i.e., ordered along one chromosome) genotype data from *5q15* of the 1000 Genomes project phase 3 [32]. The frequency of the nine ERAP1 SNPs in different populations is shown in Supplemental Table 1. Correlations between individual SNPs that indicate linkage disequilibrium are shown in Supplemental Table 2 and are generally consistent with previous studies [9]. The population frequency of the most common ERAP1 allotypes is shown in Table 1. An analogous analysis using data from the UK biobank revealed highly similar results (Supplemental Table 3) [33]. Strikingly, although 10 common allotypes constitute 99.9% of all allotypes in the European population, some populations have additional allotypes not reported before. Overall, we were able to identify at least 6 additional allotypes that have frequencies of over 0.5% in at least one population (Table 1, allotypes 11 to 16). Our analysis confirmed previous results in a larger setting, but also revealed that there is significant population variability between ERAP1 allotypes. Regardless, given the extensive use of allotypes 1-10 in the literature and their near-complete coverage of the global population (>94% globally, >99.9% in the European population) we proceeded with the functional characterization of these allotypes. Since individuals carry two copies of the ERAP1 gene, we also analyzed the combinations of allotypes present in the 2,504 samples (Figure 1A and supplemental Tables 4 and 5). The most common combination was that of allotype 8 and 2 followed by the 2-2 homozygous which cumulatively account for almost 20% of the global population. Interestingly, the combination 8-2 was found to be about twice as frequent (11%) than predicted from a random distribution (25.6% x 21.8%=5.6%). We additionally analyzed the prevalence of two SNPs in the homologous ERAP2, namely rs2549782 and rs2248374 [34, 35]. Of the four possible combinations, we were able to detect only two in our population sample, the [G,A] allotype (defined as allotype A in [35]) in 44.7% of the samples and the [T,G] allotype (defined as allotype B in [35]) in 55.3% of the samples. Since however the [G] allele of the rs2248374 leads to no detectable expression of ERAP2 [35] our results suggest that 31.3% of individuals in the database carry two [G] allotypes and should therefore have no ERAP2 protein expression. Because we previously showed that the co-occurrence of functional ERAP2 protein is dependent on the ERAP1 haplotype background [10] in Dutch and Spanish controls, we were interested to see if the ERAP1 and ERAP2 allotypes correlate in this global data set. Thus, we calculated the distribution of ERAP1 allotypes in each group of individuals carrying a particular ERAP2 allotype (Supplemental Table 6). This analysis revealed some interesting correlations. The [8,8] ERAP1 combination was found in 19.2% of individuals that carry the [A,A] ERAP2 allotype which is much more frequent compared the whole population (5.8%). In addition the [2,2] ERAP1 allotype combination was found in 22.3% of individuals carrying the [B,B] ERAP2 allotype (8.2% in the whole population). The correlation of these specific combinations of ERAP1 and ERAP2 allotypes may be functionally relevant as discussed later in this manuscript.

**Table 1:**
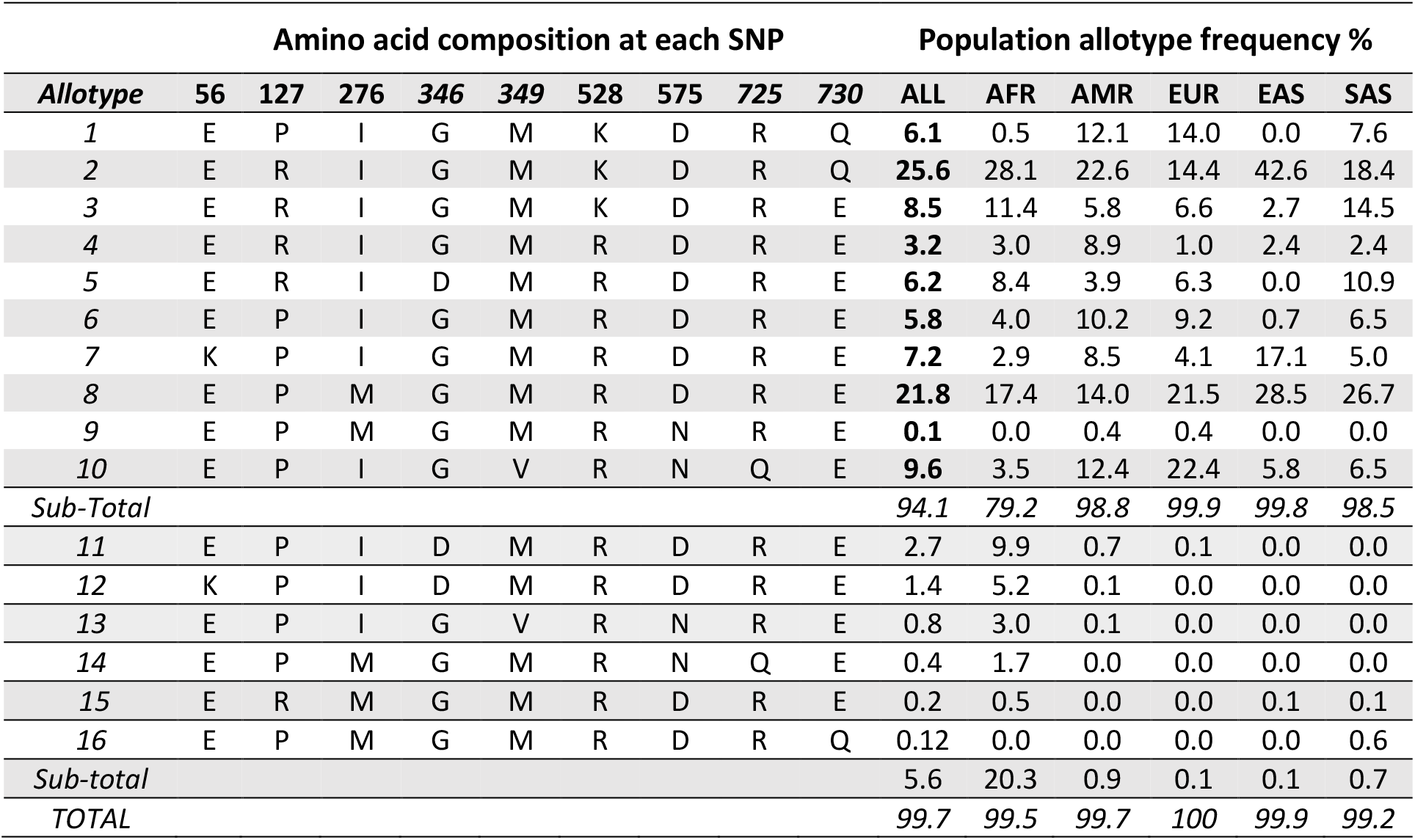
List of ERAP1 allotypes and their frequency in different populations.

| Amino acid composition at each SNP |  |  |  |  |  |  |  |  |  | Population allotype frequency % |  |  |  |  |  |
| --- | --- | --- | --- | --- | --- | --- | --- | --- | --- | --- | --- | --- | --- | --- | --- |
| Allotype | 56 | 127 | 276 | 346 | 349 | 528 | 575 | 725 | 730 | ALL | AFR | AMR | EUR | EAS | SAS |
| 1 | E | P | I | G | M | K | D | R | Q | 6.1 | 0.5 | 12.1 | 14.0 | 0.0 | 7.6 |
| 2 | E | R | I | G | M | K | D | R | Q | 25.6 | 28.1 | 22.6 | 14.4 | 42.6 | 18.4 |
| 3 | E | R | I | G | M | K | D | R | E | 8.5 | 11.4 | 5.8 | 6.6 | 2.7 | 14.5 |
| 4 | E | R | I | G | M | R | D | R | E | 3.2 | 3.0 | 8.9 | 1.0 | 2.4 | 2.4 |
| 5 | E | R | I | D | M | R | D | R | E | 6.2 | 8.4 | 3.9 | 6.3 | 0.0 | 10.9 |
| 6 | E | P | I | G | M | R | D | R | E | 5.8 | 4.0 | 10.2 | 9.2 | 0.7 | 6.5 |
| 7 | K | P | I | G | M | R | D | R | E | 7.2 | 2.9 | 8.5 | 4.1 | 17.1 | 5.0 |
| 8 | E | P | M | G | M | R | D | R | E | 21.8 | 17.4 | 14.0 | 21.5 | 28.5 | 26.7 |
| 9 | E | P | M | G | M | R | N | R | E | 0.1 | 0.0 | 0.4 | 0.4 | 0.0 | 0.0 |
| 10 | E | P | I | G | V | R | N | Q | E | 9.6 | 3.5 | 12.4 | 22.4 | 5.8 | 6.5 |
| Sub-Total |  |  |  |  |  |  |  |  |  | 94.1 | 79.2 | 98.8 | 99.9 | 99.8 | 98.5 |
| 11 | E | P | I | D | M | R | D | R | E | 2.7 | 9.9 | 0.7 | 0.1 | 0.0 | 0.0 |
| 12 | K | P | I | D | M | R | D | R | E | 1.4 | 5.2 | 0.1 | 0.0 | 0.0 | 0.0 |
| 13 | E | P | I | G | V | R | N | R | E | 0.8 | 3.0 | 0.1 | 0.0 | 0.0 | 0.0 |
| 14 | E | P | M | G | M | R | N | Q | E | 0.4 | 1.7 | 0.0 | 0.0 | 0.0 | 0.0 |
| 15 | E | R | M | G | M | R | D | R | E | 0.2 | 0.5 | 0.0 | 0.0 | 0.1 | 0.1 |
| 16 | E | P | M | G | M | R | D | R | Q | 0.12 | 0.0 | 0.0 | 0.0 | 0.0 | 0.6 |
| Sub-total |  |  |  |  |  |  |  |  |  | 5.6 | 20.3 | 0.9 | 0.1 | 0.1 | 0.7 |
| TOTAL |  |  |  |  |  |  |  |  |  | 99.7 | 99.5 | 99.7 | 100 | 99.9 | 99.2 |

**Figure 1:**
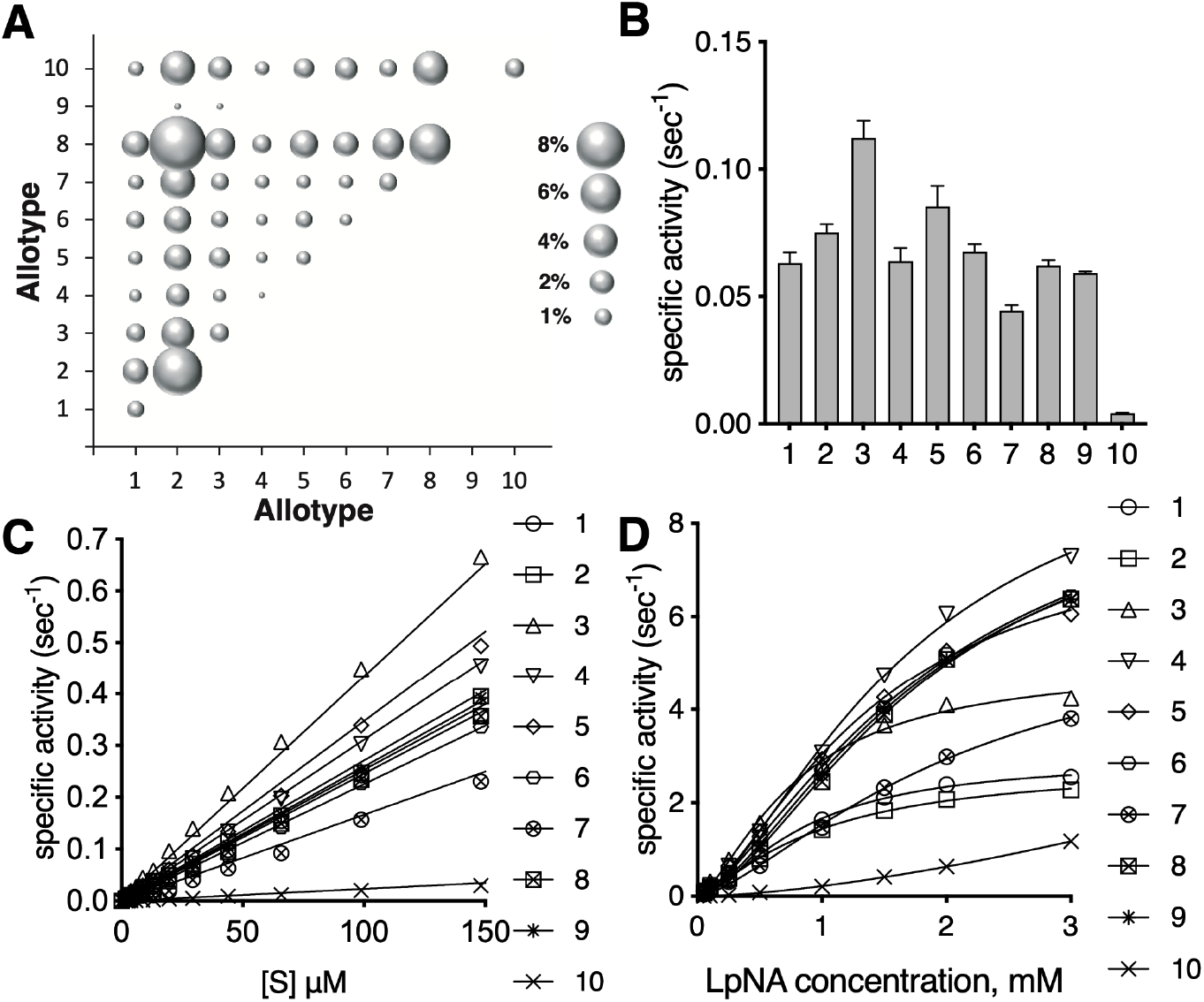
**Panel A**, bubble chart showing the relative prevalence of ERAP1 allotype combinations in the global population based on data form 1000 Genomes. **Panel B**, specific activity of ERAP1 allotypes 1-10 for the hydrolysis of dipeptidic substrate Leu-AMC. **Panel C**, specific activity of Leu-AMC hydrolysis versus substrate concentration. **Panel D**, Michaelis-Menten analysis of Leu-pNA hydrolysis by each allotype.

To characterize the enzymatic properties of the common ERAP1 allotypes, we generated recombinant ERAP1 variants corresponding to each allotype as listed in Table 1. The SNPs that define allotypes 1-10 are scattered throughout the structure of ERAP1, away from the catalytic center (Supplemental Figure 1) and can be generally categorized into two groups: a) SNPs that lie on the outside of the central cavity of the enzyme that normally accommodates the substrate (Supplemental Figure 1A) and b) SNPs that line the interior surface of this cavity and may make direct interactions with the peptide substrate (Supplemental Figure 1B).

We first characterized the ERAP1 allotypes using well-established small dipeptide substrates. The specific activity for the hydrolysis of the substrate Leu-AMC is shown in Figure 1B. There was about a 2-fold spread in specific activities for allotypes 1-9, but allotype 10 was found to be at least 10-fold less active (Figure 1B and Supplemental Figure 2). The relationship between specific activity and substrate concentration was found to be linear up to 150μM substrate which allowed the calculation of the k_cat_/K_M_ ratio for Leu-AMC (Figure 1C and Supplemental Table 7). Allotype 3 was found to be the most active of all and allotype 10 was 18-fold less active. To obtain full Michaelis-Menten analysis we employed the chromogenic dipeptide substrate Leu-pNA [36]. Data fit best to an allosteric MM model as previously demonstrated [28, 36] allowing us to calculate the enzymatic parameters (Figure 1D and Supplemental Table 7). This analysis demonstrated that the changes in specific activity between allotypes are due both to changes in affinity for the substrate (K_half_) and to changes in maximal catalytic efficiency (V_max_). Similar to Leu-AMC, allotype 10 was much less active in hydrolyzing Leu-pNA, which unfortunately precluded reliable calculation of K_half_ and V_max_ for this allotype.

**Figure 2:**
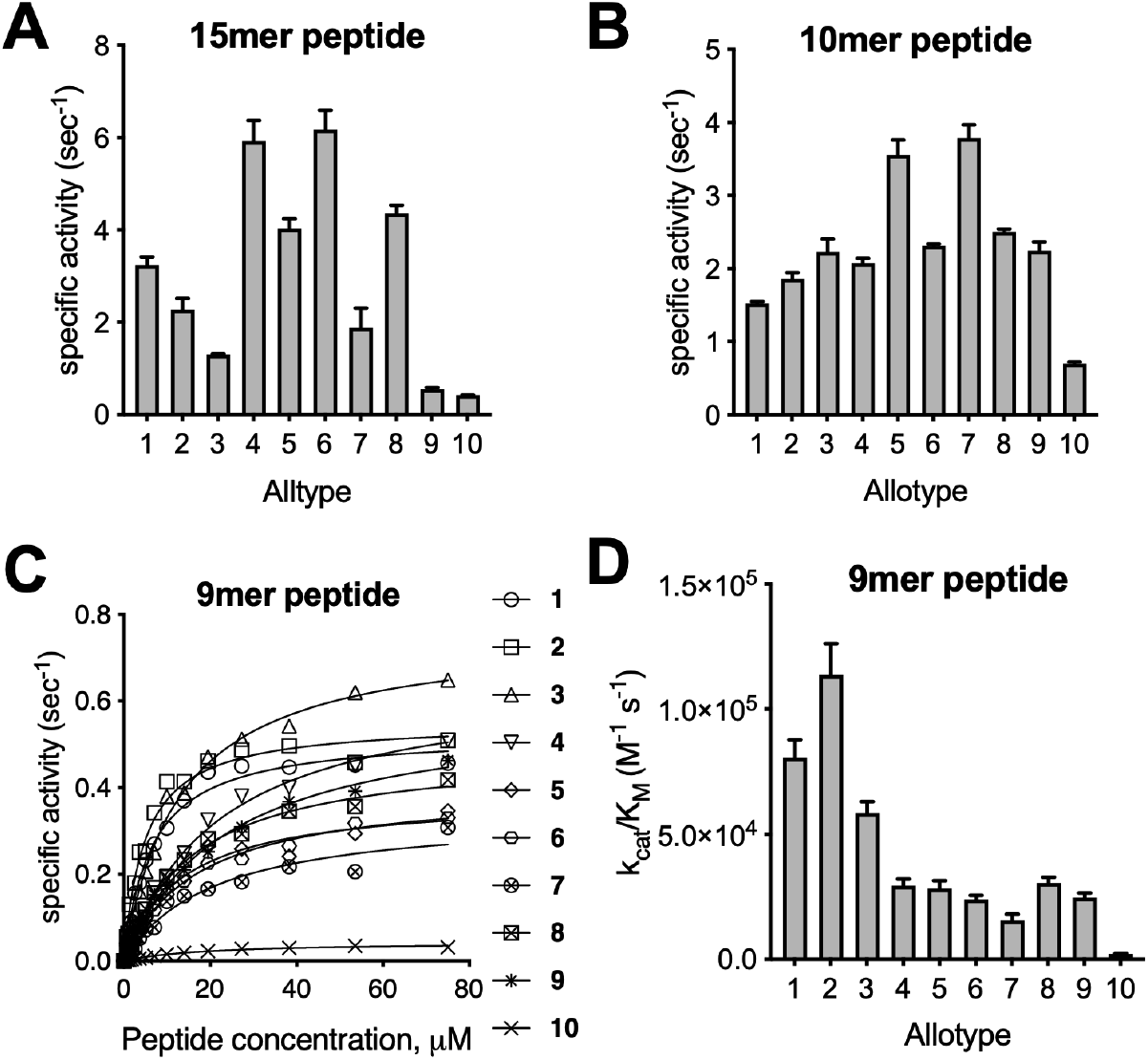
Hydrolysis of peptidic substrates by ERAP1 allotypes. **Panel A**, specific activity of the hydrolysis of 15mer peptide with the sequence LLRIQRGPGRAFVTI. **Panel B**, specific activity of the hydrolysis of 10mer peptide with the sequence LLKHHAFSFK. **Panel C**, Michaelis-Menten analysis of the hydrolysis of the 9mer peptide YTAFTIPSI by ERAP1 allotypes. **Panel D**, catalytic efficiency (k_cat_/K_M_) of each ERAP1 allotype for the trimming of the 9mer epitope YTAFTIPSI.

Since ERAP1 trims long N-terminally extended peptide precursors of antigenic peptides, we turned to more physiologically-relevant long peptides. Recently determined co-crystal structures of ERAP1 with bound 15mer and 10mer peptide analogues revealed that the peptides are processed in a large internal cavity, while making interactions with residues that line that cavity, which can drive selectivity [16]. We measured the rate of N-terminus hydrolysis of two peptides of a similar backbone sequence as the co-crystalized analogues, namely the 15mer LLRIQRGPGRAFVTI and the 10mer LLKHHAFSFK (Figure 2A and 2B). Similar to the results with the small substrates we recorded a significant variation in trimming rates. Although allotype 10 was again the least efficient, processing rate differences were less pronounced compared to smaller substrates. Notably, allotype 10 was about as efficient as allotype 9 in trimming the 15mer. Interestingly, the pattern between the two peptides was significantly different and while allotypes 4 and 6 trimmed the 15mer the fastest, allotypes 5 and 7 trimmed the 10mer the fastest. This results are consistent with a complex landscape of peptide-enzyme interactions that drive specificity as previously proposed and suggest that the effect of allotype variation may be substrate-dependent [16, 37].

To better understand the mechanism behind the variation of trimming rates, we utilized a recently developed assay suitable for Michaelis-Menten analysis that follows the trimming of a 9mer antigenic epitope with the sequence YTAFTIPSI (Figure 2C) [30]. Using this assay we calculated k_cat_ and K_M_ for each ERAP1 allotype (Supplemental Table 8). The turnover rate (k_cat_), a measure of the maximal catalytic rate of hydrolysis, varied by only about 2-fold between allotypes 1-9, but was about 10-fold reduced for allotype 10, indicating that this allotype is catalytically deficient. The Michaelis constant (K_M_), a measure of how well the enzyme can recognize the substrate, varied between allotypes by up to 6-fold with allotypes 1 and 2 having the highest affinity for the substrate. The ratio k_cat_/K_M_, a measure of the overall catalytic efficiency of the enzyme, varied up to 60-fold, indicating that changes in k_cat_ and K_M_ can synergize to enhance differences in trimming rates for particular substrates (Figure 2D). Notably, allotype 10 had a 60-fold lower catalytic efficiency compared to allotype 2 because of the combined effect of lower substrate affinity and lower turnover rate. Thus, we conclude that polymorphic variation in ERAP1 can influence peptide trimming rates by affecting both catalytic efficiency and substrate recognition.

ERAP1 trimming of antigenic epitope precursors in the ER often includes multiple trimming steps and which intermediates accumulate can affect which peptides will bind onto MHCI[38]. To evaluate the effect of ERAP1 allotype variation on sequential trimming reactions we followed the generation of the ovalbumin epitope SIINFEKL from the starting 14mer extended epitope GLEQLE<u>SIINFEKL</u> (Figure 3). In all cases the 14mer was catabolized and the 8mer produced but with variable efficiencies. As before, allotype 10 was less active in trimming and it was necessary to use at a 10-fold higher concentration to properly follow the reaction. In all reactions, all possible intermediates were detected (Figure 3A). However, we observed significant differences both between intermediate accumulation and on the overall rate of mature epitope generation (Figure 3B). Specifically, allotypes 1-3 accumulated the mature epitope the fastest, whereas allotypes 5 and 7 were less efficient, in part because of slower trimming of the initial 14mer. This comes as a sharp contrast to their ability to trim the 10mer peptide shown in Figure 3B. Interestingly, allotype 5 accumulated significant amounts of the 11mer intermediate. Several other allotypes (4 and 6-10) accumulated the 12mer intermediate. Since peptides of 10-12 residues can be immunogenic, accumulation of distinct intermediates by different allotypes could contribute to differences in immune responses between individuals.

**Figure 3:**
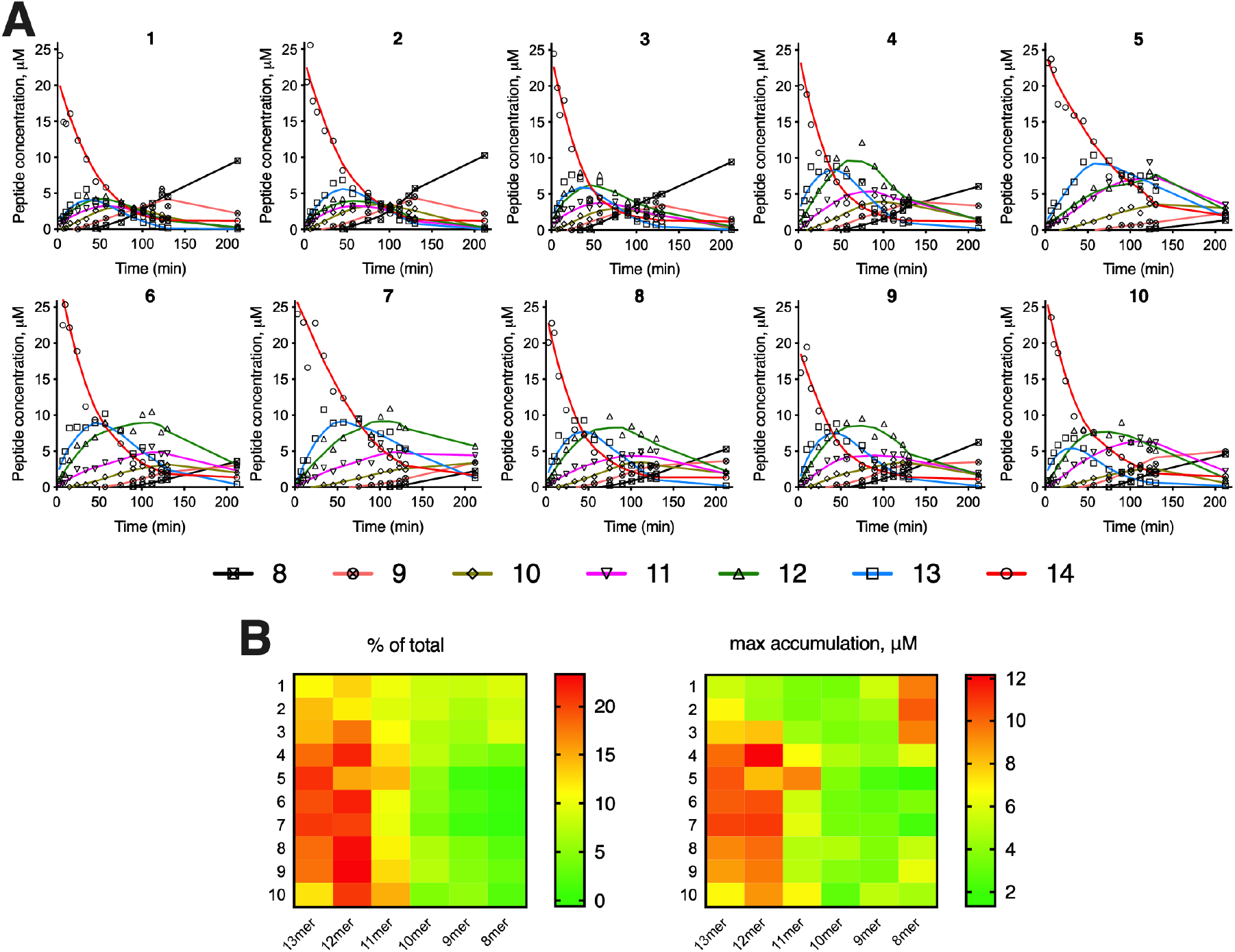
**Panel A**, trimming of the 14mer peptide GLEQLESIINFEKL, precursor to the ovalbumin epitope SIINFEKL, by ERAP1 allotypes. Each peptide is quantified by Mass Spectrometry and its concentration is plotted as a function of reaction time. Lines correspond to locally weighted scatterplot smoothing (LOWESS) and are only used as a visual aid to follow peptide accumulation. Allotype 10 was used at a 10-fold higher concentration. **Panel B**, left: heatmap indicating accumulation of each peptide intermediate as a fraction of the total amount of peptides in the reaction; right: heatmap showing the maximal accumulation of each peptide intermediate during the reaction.

ERAP1 is an emerging pharmacological target for cancer immunotherapy and the control of inflammatory autoimmunity, including rheumatic conditions such as ankylosing spondylitis [39, 40]. Given the wide distribution of common allotypes in the population, it is crucial to know if inhibitors with clinical potential can effectively inhibit all allotypes. We performed inhibition titrations using Leu-AMC and inhibitors DG013A and GSK849, both shown to be active in cellular assays. DG013A is a potent transition state analogue that targets the active site of the enzyme [26]. GSK849 targets a regulatory site of ERAP1 and while is an activator for small substrates, it inhibits long peptide hydrolysis by interfering with C-terminus recognition [30]. DG013A was able to inhibit all 10 allotypes with high potency (pIC_50_ between 7.2 and 7.6) (Figure 4A). GSK849 acted as an apparent activator of small substrate hydrolysis as previously reported [30] and its efficacy varied significantly between allotypes (pXC_50_ between 4.8 and 6.5)(Figure 4B). GSK849 was, however, an inhibitor of the more physiologically-relevant 9mer substrate YTAFTIPSI (Figure 4C). A comparison of the pIC_50_ and pXC_50_ values for the two substrates is shown in Figure 4D. GSK849 was most active against allotypes 1 and 2 and least active versus allotype 10. Overall, there was a positive correlation between allotype activity and GSK849’s ability to inhibit (Figure 4E). Additionally, there was a positive correlation between the pXC_50_ value and the fractional activation observed (Figure 4F). This surprising finding suggests that the regulatory, but not the catalytic site of ERAP1, is sensitive to the allotypic state of the enzyme.

**Figure 4:**
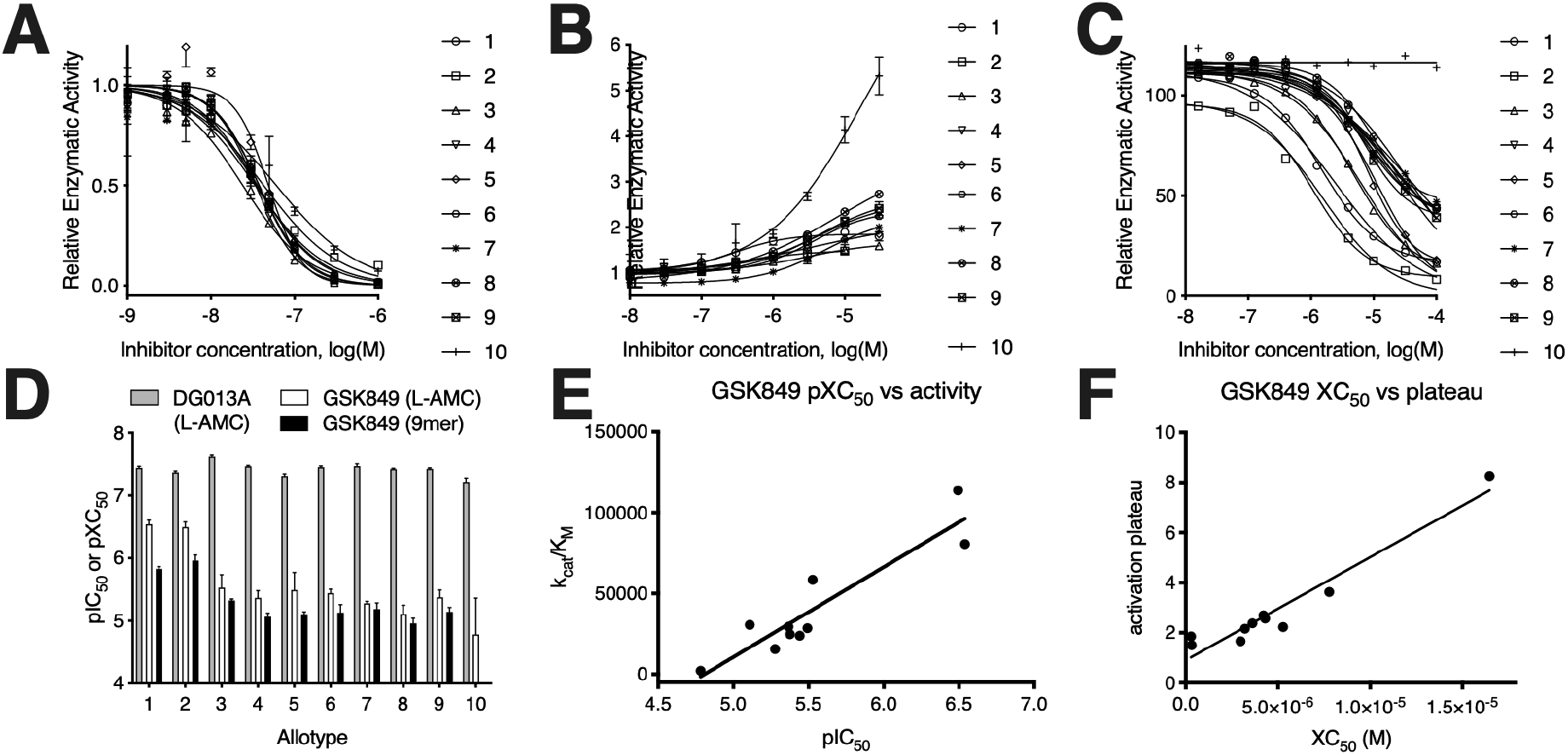
Effect of inhibitors DG013A and GSK849 on the activity of ERAP1 allotypes. **Panel A**, effect of titration of DG013A on the hydrolysis of Leu-AMC by ERAP1 allotypes. **Panel B**, effect of titration of GSK849 on the hydrolysis of Leu-AMC. **Panel C**, ability of GSK849 to inhibit hydrolysis of a 9mer peptide by ERAP1 allotypes. **Panel D**, comparison of pIC_50_ or pXC_50_ values from the titrations shown in panels A-C. **Panel E**, correlation between allotype specific activity and pIC_50_ value for GSK849. **Panel F**, correlation between maximal activity enhancement by GSK849 and XC_50_ of each allotype.

The high variability in enzymatic activity between ERAP1 allotypes suggests that individuals carrying different combinations of allotypes can have an even larger variability of ERAP1 activities. Assuming that no specific interactions exist between the two alleles, the total enzymatic activity of an individual would be expected to be the sum of the activities of each allele. A caveat in this analysis is that SNPs and therefore allotypes may affect gene expression or protein turnover, thus affecting the steady-state protein levels. Indeed, two recent reports suggested that SNPs can affect ERAP1 expression to some degree something that could either exacerbate or ameliorate allotype activity variation [18, 19]. Since however, existing studies were limited to effects of specific SNPs and not allotypes and effects on expression levels were relatively small, the potential effect of altered levels of expression are not examined here. To calculate the expected total activity of the two alleles carried by individuals we utilized the measurements of activity for the 9mer substrate since we had obtained reliable catalytic efficiency measurements (k_cat_/K_M_, Figure 2D and Supplemental Table 8). A plot of expected specific activity versus population frequency is shown in Figure 5A. A bubble chart showing the population frequency and expected enzymatic activity for each possible combination of ERAP1 allotype is shown in Figure 5B. Individuals carrying different common combinations of allotypes are expected to possess a wide range, of about 60-fold, of total ERAP1 activity. Most common allotype combinations fall within a more limited range, about 10-fold (cyan region, Figure 5A). Homozygous individuals of allotype 2 are quite frequent in the global population and would be expected to feature the highest ERAP1 activity (red region, Figure 5A). Combinations of 2 with 8 are also very common and have moderate-to-high enzyme activity (magenta region, Figure 5A). Several moderately-active combinations of allotype 9 are rare in the population, as is the [4,4] homozygous. Finally, homozygous individuals of allotype 10 are found in ∼1.2% of the global population and should have the lowest ERAP1 activity, being functional knockouts for some peptide substrates. Significant variability in both activity and frequency distribution was found in different populations (Supplemental Figure 3), a phenomenon that may signify local host-pathogen balancing selection pressures, a notion previously suggested for individual SNPs [41].

**Figure 5:**
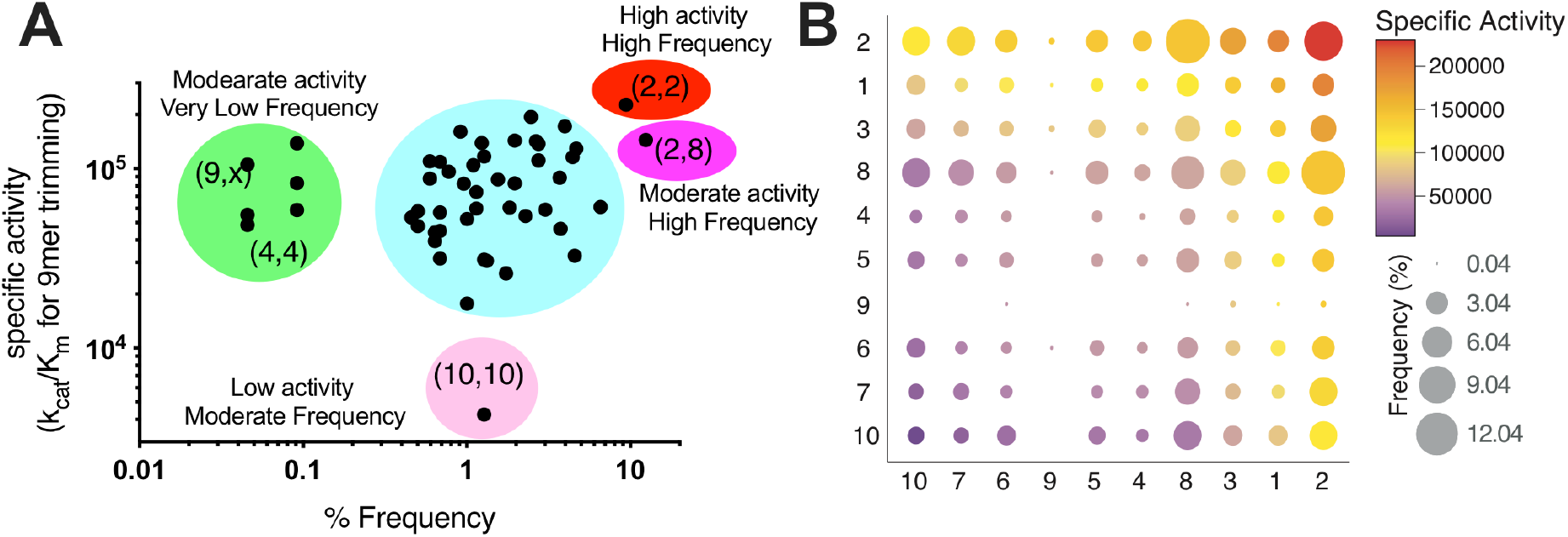
The landscape of ERAP1 activities in the population. **Panel A**, scatterplot showing the correlation between total estimated ERAP1 activity and frequency of allotype combinations found in individuals. Specific allotype combinations are indicated in parenthesis. **Panel B**, bubble chart showing the frequency of allotype combinations color-coded by their estimated activity. Allotypes have been clustered based on activity (highest activity top right, lowest bottom left).

## Discussion

A wealth of genetic association studies has linked ERAP1 polymorphic variation to the existence of coding SNPs [7, 42] which have been reported to affect enzyme function, thus providing mechanistic support for observed epistasis between ERAP1 and HLA [14, 15, 22, 34]. In general, functional effects of single SNPs are relatively modest but complex and hint at possible synergism between SNPs that is poorly understood. This is highlighted in cases in which common allotypes harbor SNPs that protect along with SNPs that predispose to disease [21]. Thus, a deep understanding of the complex patterns of disease susceptibility observed in genetic-association studies requires a detailed knowledge of the functional differences between naturally-occurring allotypes and not just SNPs. Our results here suggest that SNPs can synergize to affect function in a substrate-dependent manner. This is probably best highlighted for allotype 10 which is up to 60-fold less active than other allotypes. Such a strong functional change has not been reported before for individual SNPs in ERAP1 and is probably the result of synergism. Specifically, allotype 10 carries the SNP 528R that has been reported to associate with decreased risk to Ankylosing Spondylitis and Psoriasis and to reduce enzyme activity by about 2-fold [13, 14, 20, 43]. In addition, it carries the SNP 575N which has been reported to affect enzymatic activity depending on the polymorphic state of position 528 [44]. These two SNPs may synergize along with two additional SNPs unique to this allotype: i) 349V, that is relatively close to the catalytic site although it is a conservative alteration that has not been reported to affect activity in isolation and ii) 725Q, that has been demonstrated to reduce enzymatic activity [45] and lies in the interface between domains II and IV and could induce changes in the dynamics of ERAP1’s conformational change, similar to position 528 [46]. It is possible that synergism between those four SNPs could underlie the greatly reduced catalytic efficiency of allotype 10. Strikingly, this reduction appears to be partially substrate-dependent. Trimming of the 10mer substrate was less affected by this allotype compared to the other substrates we tested. This could be due to unproductive interactions between the C-terminal side-chain of this peptide and 725R as observed in a recent crystal structure, making the substitution 725Q favorable for this particular substrate [16]. This observation highlights a motif that can be seen throughout our results. While some changes in activity between allotypes are consistent, they can be influenced by the substrate. This is remarkably reminiscent of the effects of polymorphic variation in MHC molecules. Changes in the shape and dynamics of the peptide binding groove, affect the binding of different peptides both thermodynamically and kinetically and contribute to the variability of immune responses [47]. It appears that ERAP1 allotypes operate in the same general principle as MHC haplotypes. They induce variability of substrate processing, in a substrate-dependent manner, thus contributing, in tandem with MHC polymorphic variation, to immune response variability within the population.

While it is difficult to dissect the mechanism that underlies the role of each SNP to the activity of each allotype without additional structural information, some insight can still be extracted. Previous studies demonstrated that 528K results in higher enzymatic activity possibly due to effects on the conformational dynamics of the enzyme [46]. Accordingly, allotypes 1-3 which all carry this SNP, are amongst the most active in our assays. The polymorphism 730Q has been suggested to affect activity due to changes in interaction with the C-terminal moiety of the peptide [7, 46]. Accordingly, allotypes 1 and 2, which carry this SNP are more active versus the 9mer substrate that carries a hydrophobic C-terminal side-chain; allotypes 3-10 that have 730E at that location, a negative charge, would be expected to be worse in interacting with this substrate, all are less active compared to 1 and 2. Furthermore, allotypes that carry the 127P polymorphism (also reported to be associated with AS [21]) tend to have lower activity, although no specific effects of this SNP have been reported before [7]. This polymorphic location lies on a putative substrate exit-channel and the reduced structural flexibility in the mouth of this cannel due to the proline residue may reduce the kinetics of product-substrate exchange leading to a slower apparent activity [16]. Finally, the sensitivity of the regulatory site to inhibition appears to be allotype dependent, a notion that is consistent with a previous study from our group that suggested that the regulatory site communicates with the active site though effects on the conformational dynamics of the enzyme [28]. Further structural analysis of these common ERAP1 allotypes will be invaluable in dissecting synergism between SNPs that underlie functional changes in allotypes.

Although ERAP2 has been described as an accessory aminopeptidase that supplements ERAP1 activity, recent genetic association data have pointed to important roles in both cancer immunotherapy and autoimmunity [48-50]. Because the rs2248374 polymorphism that leads to lack of protein expression [35], the effects on functional ERAP2 activity in the cell can be pronounced since individuals homozygous for the G allele express no full-length enzymatically-active ERAP2 and heterozygous individuals express half the canonical protein amount. Although ERAP2 has different specificity than ERAP1 [51, 52], since there are the only known ER-resident aminopeptidases, they cumulatively define the spectrum of aminopeptidase activity in the ER. From the genetic and enzymatic analysis, it appears that the highest ERAP2 expression allotype combination is found more frequently in individuals that carry an intermediate activity ERAP1 allotype (allotype 8), whereas individuals with reduced ERAP2 protein expression, are more often homozygous for the most active ERAP1 allotype (allotype 2). This is in line with previous genetic analysis of co-occurrence of ERAP1-ERAP2 haplotypes [10]. Thus, it appears that some balancing selection may exist that attempts to normalize total aminopeptidase activity in the ER.

Although our *in vitro* analysis has the advantage of allowing the accurate determination of fundamental molecular properties, how these properties translate to changes in antigen presentation by cells is not always straightforward. Antigen presentation is primarily controlled by peptide binding onto MHCI and this process may mask some changes in ERAP1 activity. Indeed, recent studies have suggested that the potential of ERAP1 in influencing antigen presentation is limited by this phenomenon [53, 54]. Still, many recent cell-based studies using proteomic approaches have provided strong support that changes in ERAP1 activity due to polymorphic variation are translated to changes in antigen presentation [55]. Also, although the expression of ERAP1 mediated by splice interfering variants [18] is in linkage disequilibrium with variants that encode distinct allotypes, genetic association studies have shown that for some (HLA-associated) conditions the disease risk is primarily mapped to enzymatic activity [10]. Thus, although MHCI binding is the dominant filter, changes in ERAP1 activity are highly relevant to antigen presentation and probably underlie part of known disease associations.

In summary, we provide a current allotype and genotype analysis for the ERAP1 gene in human populations and a detailed enzymatic characterization of the 10 most common ERAP1 allotypes. Our results suggest that individual SNPs synergize to shape allotype enzymatic properties by affecting both catalytic efficiency and substrate affinity. Our analysis defines a two-order of magnitude-wide landscape of ERAP1 genotype activities in human populations and suggests that ERAP1/2 genotypes operate in tandem with MHCI haplotypes to generate the necessary plurality in antigen presentation that supports the observed variability of immune responses between individuals.

## ACKNOWLEDGMENTS

This research has been funded in part by internal funds of the National Centre for Scientific Research “Demokritos” and the UK Biobank Resource (application no. 20361). Jonas J. W. Kuiper is supported by a VENI award from the Netherlands Organization for Scientific Research (N.W.O. project number 016.186.006).

