## Supplemental Data for "Common allotypes of ER aminopeptidase 1 have substrate-dependent and highly variable enzymatic properties"

**Affiliations:**
[1]Medicinal Science and Technology, GlaxoSmithKline, Stevenage, Hertfordshire SG1 2NY, U.K.
[2]National Centre for Scientific Research "Demokritos", Athens 15341, Greece.
[3]Department of Ophthalmology, University Medical Center Utrecht, Utrecht University, Utrecht, The Netherlands. Center for Translational Immunology, University Medical Center Utrecht, University of Utrecht, Utrecht, Netherlands.
[4] Human Genetics, GlaxoSmithKline, Stevenage, Hertfordshire SG1 2NY, U.K.
[5] Adaptive Immunity Research Unit, GlaxoSmithKline, Stevenage, Hertfordshire SG1 2NY, U.K.
[6] Laboratory of Biochemistry, Department of Chemistry, National and Kapodistrian University of Athens, Panepistimiopolis Zografou 157 84, Greece.

[#] These authors contributed equally to the manuscript
[*].

**Supplemental Table 1:** Frequency of ERAP1 SNPs in different populations based on analysis of 2,504 Human Genomes from the 1000 Genomes Project.

| SNP ID / Amino acid | Population Frequency of SNP % | | | | | | | | | | | |
|---|---|---|---|---|---|---|---|---|---|---|---|---|
| | ALL | | AFR | | AMR | | EUR | | EAS | | SAS | |
| rs3734016 | C | T | C | T | C | T | C | T | C | T | C | T |
| E56K | 91.4 | 8.6 | 91.9 | 8.1 | 91.4 | 8.6 | 95.9 | 4.1 | 82.8 | 17.2 | 95.0 | 5.0 |
| rs26653 | G | C | G | C | G | C | G | C | G | C | G | C |
| P127R | 56.3 | 43.8 | 48.2 | 51.8 | 58.8 | 41.2 | 71.8 | 28.2 | 52.1 | 47.9 | 53.7 | 46.3 |
| rs26618 | T | C | T | C | T | C | T | C | T | C | T | C |
| I276M | 77.3 | 22.7 | 80.4 | 19.6 | 85.6 | 14.4 | 78.1 | 21.9 | 71.4 | 34.6 | 72.6 | 27.4 |
| rs27895 | C | T | C | T | C | T | C | T | C | T | C | T |
| G346D | 89.6 | 10.4 | 76.2 | 23.8 | 95.2 | 4.8 | 93.6 | 6.4 | 99.9 | 0.1 | 89.1 | 10.9 |
| rs2287987 | T | C | T | C | T | C | T | C | T | C | T | C |
| M349V | 89.5 | 10.5 | 93.4 | 6.6 | 87.3 | 12.7 | 77.5 | 22.5 | 94.2 | 5.8 | 93.1 | 6.9 |
| rs30187 | C | T | T | C | C | T | T | C | C | T | C | T |
| K528R | 59.6 | 40.4 | 40.2 | 59.8 | 59.5 | 40.5 | 35.0 | 65.0 | 54.7 | 45.3 | 59.1 | 40.9 |
| rs10050860 | C | T | C | T | C | T | C | T | C | T | C | T |
| D575N | 89.4 | 10.6 | 93.5 | 6.5 | 86.9 | 13.1 | 77.1 | 22.9 | 94.2 | 5.8 | 93.1 | 6.9 |
| rs17482078 | C | T | C | T | C | T | C | T | C | T | C | T |
| R725Q | 89.9 | 10.1 | 94.6 | 5.4 | 87.6 | 12.4 | 77.6 | 22.4 | 94.2 | 5.8 | 93.5 | 6.5 |
| rs27044 | G | C | G | C | G | C | G | C | G | C | G | C |
| Q730E | 32.1 | 67.9 | 28.9 | 71.1 | 34.9 | 65.1 | 28.5 | 71.5 | 42.9 | 57.1 | 26.9 | 73.1 |

**Supplemental Table 2:** Correlation of coding SNPs in the 1000 Genomes Project data set suggesting linkage disequilibrium between particular SNPs. Number correspond to percentage of frequency of the two SNPs exist in an individual. Color coding indicates level of correlation (green=0, red=100).

| | 56E | 56K | 127R | 127P | 276I | 276M | 346G | 346D | 349M | 349V | 528K | 528R | 575D | 575N | 725R | 725Q |
|---|---|---|---|---|---|---|---|---|---|---|---|---|---|---|---|---|
| **127R** | 43.8 | 0.0 | | | | | | | | | | | | | | |
| **127P** | 47.7 | 8.6 | | | | | | | | | | | | | | |
| **276I** | 68.8 | 8.6 | 43.6 | 33.8 | | | | | | | | | | | | |
| **276M** | 22.7 | 0.0 | 0.2 | 22.5 | | | | | | | | | | | | |
| **346G** | 82.5 | 7.2 | 37.5 | 52.1 | 67.0 | 22.7 | | | | | | | | | | |
| **346D** | 9.0 | 1.4 | 6.2 | 4.2 | 10.4 | 0.0 | | | | | | | | | | |
| **349M** | 80.9 | 8.6 | 43.7 | 45.8 | 66.8 | 22.7 | 79.1 | 10.4 | | | | | | | | |
| **349V** | 10.5 | 0.0 | 0.0 | 10.5 | 10.5 | 0.0 | 10.5 | 0.0 | | | | | | | | |
| **528K** | 40.4 | 0.0 | 34.2 | 6.2 | 40.4 | 0.0 | 40.3 | 0.0 | 40.4 | 0.0 | | | | | | |
| **528R** | 51.0 | 8.6 | 9.6 | 50.1 | 37.0 | 22.7 | 49.3 | 10.3 | 49.1 | 10.5 | | | | | | |
| **575D** | 80.8 | 8.6 | 43.7 | 45.7 | 66.9 | 22.5 | 79.0 | 10.4 | 89.4 | 0.0 | 40.4 | 49.0 | | | | |
| **575N** | 10.6 | 0.0 | 0.0 | 10.6 | 10.5 | 0.1 | 10.6 | 0.0 | 0.1 | 10.5 | 0.0 | 10.6 | | | | |
| **725R** | 81.4 | 8.6 | 43.7 | 46.7 | 67.7 | 22.2 | 79.6 | 10.4 | 89.0 | 9.6 | 40.3 | 49.6 | 88.9 | 1.0 | | |
| **725Q** | 10.1 | 0.0 | 0.1 | 10.0 | 9.6 | 0.4 | 10.1 | 0.0 | 0.5 | 0.9 | 0.0 | 10.0 | 0.5 | 9.6 | | |
| **730Q** | 32.1 | 0.0 | 25.7 | 6.4 | 32.0 | 0.1 | 32.0 | 0.1 | 32.0 | 0.1 | 31.8 | 0.3 | 32.0 | 0.1 | 32.1 | 0.0 |
| **730E** | 59.4 | 8.6 | 18.1 | 49.9 | 45.4 | 22.5 | 57.7 | 10.3 | 57.6 | 10.4 | 8.6 | 59.3 | 57.4 | 10.5 | 57.9 | 10.1 |

**Supplemental Table 3:** Frequency of ERAP1 allotypes in samples from the UK biobank stratified by self-reported ethnic background.

| | ***Population allotype frequency %*** | | | | | | | | | | | | |
|---|---|---|---|---|---|---|---|---|---|---|---|---|---|
| ***Allotype*** | **British** | **Irish** | **Any other white background** | **White and Black Caribbean** | **White and Black African** | **White and Asian** | **Any other mixed background** | **Indian** | **Bangladeshi** | **Any other Asian background** | **Caribbean** | **African** | **Any other Black background** |
| *1* | 13.0 | 13.0 | 13.9 | 8.4 | 9.0 | 10.2 | 9.4 | 8.2 | 8.6 | 10.9 | 6.7 | 2.5 | 0.7 |
| *2* | 13.6 | 12.6 | 14.0 | 18.8 | 21.0 | 19.6 | 20.2 | 20.6 | 19.0 | 23.3 | 25.0 | 26.6 | 28.7 |
| *3* | 7.0 | 6.8 | 8.4 | 10.0 | 8.8 | 9.9 | 9.6 | 13.6 | 13.3 | 13.1 | 12.0 | 10.8 | 12.1 |
| *4* | 0.6 | 0.5 | 0.8 | 1.6 | 1.4 | 1.1 | 1.8 | 1.5 | 1.9 | 2.5 | 1.8 | 2.6 | 2.8 |
| *5* | 6.7 | 6.8 | 6.9 | 6.4 | 9.2 | 7.1 | 6.2 | 11.1 | 12.2 | 8.4 | 8.2 | 7.7 | 8.5 |
| *6* | 8.5 | 7.6 | 9.2 | 7.6 | 6.2 | 8.5 | 7.6 | 7.9 | 7.6 | 5.4 | 6.8 | 5.7 | 5.7 |
| *7* | 4.5 | 5.9 | 3.3 | 3.6 | 2.9 | 4.6 | 4.7 | 4.9 | 4.0 | 5.2 | 6.2 | 2.4 | 2.2 |
| *8* | 22.9 | 22.2 | 23.6 | 20.4 | 18.7 | 22.4 | 22.7 | 21.3 | 22.2 | 21.0 | 24.3 | 17.5 | 16.4 |
| *9* | 0.2 | 0.5 | 0.1 | 0.2 | 0.0 | 0.2 | 0.1 | 0.0 | 0.0 | 0.0 | 0.0 | 0.0 | 0.0 |
| *10* | 22.0 | 23.1 | 18.9 | 14.8 | 13.2 | 15.3 | 14.3 | 9.5 | 9.9 | 8.6 | 8.2 | 5.3 | 3.7 |

**Supplemental Table 4:** Distribution of combinations of ERAP1 allotypes in the global population

| *% in population (global)* | | | | | | | | | | |
|---|---|---|---|---|---|---|---|---|---|---|
| ***Allotype*** | ***1*** | ***2*** | ***3*** | ***4*** | ***5*** | ***6*** | ***7*** | ***8*** | ***9*** | ***10*** |
| ***1*** | 0.9 | | | | | | | | | |
| ***2*** | 2.2 | 8.2 | | | | | | | | |
| ***3*** | 1.1 | 3.5 | 1.1 | | | | | | | |
| ***4*** | 0.5 | 1.7 | 0.5 | 0.1 | | | | | | |
| ***5*** | 0.6 | 2.4 | 1.4 | 0.4 | 0.6 | | | | | |
| ***6*** | 1.0 | 2.4 | 0.9 | 0.4 | 0.9 | 0.4 | | | | |
| ***7*** | 0.7 | 4.1 | 1.0 | 0.6 | 0.6 | 0.6 | 1.1 | | | |
| ***8*** | 2.4 | 11.0 | 3.2 | 1.1 | 2.6 | 2.0 | 3.3 | 5.8 | | |
| ***9*** | 0.0 | 0.1 | 0.1 | 0.0 | 0.0 | 0.0 | 0.0 | 0.0 | 0.0 | |
| ***10*** | 1.7 | 4.0 | 1.8 | 0.6 | 1.3 | 1.6 | 1.0 | 4.1 | 0.0 | 1.2 |

**Supplemental Table 5:** Distribution of combinations of ERAP1 allotypes in the sub-populations

| ***% in population (EUR)*** | | | | | | | | | | |
|---|---|---|---|---|---|---|---|---|---|---|
| ***Allotype*** | ***1*** | ***2*** | ***3*** | ***4*** | ***5*** | ***6*** | ***7*** | ***8*** | ***9*** | ***10*** |
| ***1*** | 3.0 | | | | | | | | | |
| ***2*** | 3.6 | 2.4 | | | | | | | | |
| ***3*** | 2.0 | 1.0 | 1.0 | | | | | | | |
| ***4*** | 0.4 | 0.2 | 0.0 | 0.0 | | | | | | |
| ***5*** | 1.6 | 1.8 | 0.4 | 0.0 | 0.6 | | | | | |
| ***6*** | 2.0 | 4.0 | 1.2 | 0.0 | 1.4 | 0.8 | | | | |
| ***7*** | 1.6 | 0.8 | 0.2 | 0.0 | 0.6 | 0.8 | 0.2 | | | |
| ***8*** | 6.2 | 5.6 | 1.8 | 1.0 | 2.4 | 2.6 | 2.0 | 6.2 | | |
| ***9*** | 0.2 | 0.2 | 0.2 | 0.0 | 0.0 | 0.2 | 0.0 | 0.0 | 0.0 | |
| ***10*** | 4.6 | 6.8 | 4.4 | 0.4 | 3.2 | 4.8 | 1.8 | 9.2 | 0.0 | 4.8 |

| ***% in population (AMR)*** | | | | | | | | | | |
|---|---|---|---|---|---|---|---|---|---|---|
| ***Allotype*** | ***1*** | ***2*** | ***3*** | ***4*** | ***5*** | ***6*** | ***7*** | ***8*** | ***9*** | ***10*** |
| ***1*** | 1.4 | | | | | | | | | |
| ***2*** | 5.8 | 5.2 | | | | | | | | |
| ***3*** | 1.4 | 2.3 | 1.4 | | | | | | | |
| ***4*** | 2.0 | 4.6 | 1.4 | 0.6 | | | | | | |
| ***5*** | 0.3 | 1.4 | 0.6 | 0.3 | 0.6 | | | | | |
| ***6*** | 2.3 | 4.6 | 0.9 | 2.0 | 0.3 | 1.2 | | | | |
| ***7*** | 1.2 | 3.5 | 0.3 | 2.3 | 0.9 | 1.4 | 2.3 | | | |
| ***8*** | 3.5 | 6.6 | 0.9 | 2.3 | 1.7 | 2.9 | 1.2 | 2.3 | | |
| ***9*** | 0.0 | 0.3 | 0.3 | 0.0 | 0.0 | 0.0 | 0.0 | 0.3 | 0.0 | |
| ***10*** | 4.6 | 4.9 | 0.6 | 1.7 | 0.9 | 3.7 | 1.7 | 4.0 | 0.0 | 0.9 |

| ***% in population (AFR)*** | | | | | | | | | | |
|---|---|---|---|---|---|---|---|---|---|---|
| ***Allotype*** | ***1*** | ***2*** | ***3*** | ***4*** | ***5*** | ***6*** | ***7*** | ***8*** | ***9*** | ***10*** |
| ***1*** | 0.0 | | | | | | | | | |
| ***2*** | 0.2 | 8.6 | | | | | | | | |
| ***3*** | 0.2 | 6.5 | 0.9 | | | | | | | |
| ***4*** | 0.0 | 2.0 | 0.8 | 0.0 | | | | | | |
| ***5*** | 0.2 | 4.2 | 2.1 | 0.9 | 0.3 | | | | | |
| ***6*** | 0.2 | 1.7 | 0.8 | 0.3 | 1.1 | 0.2 | | | | |
| ***7*** | 0.2 | 1.7 | 1.2 | 0.5 | 0.2 | 0.2 | 0.2 | | | |
| ***8*** | 0.2 | 8.9 | 3.3 | 0.5 | 3.5 | 1.4 | 0.9 | 3.6 | | |
| ***9*** | 0.0 | 0.0 | 0.0 | 0.0 | 0.0 | 0.0 | 0.0 | 0.0 | 0.0 | |
| ***10*** | 0.0 | 2.3 | 1.2 | 0.0 | 0.9 | 0.2 | 0.2 | 1.1 | 0.0 | 0.0 |

| % in population (EAS) | | | | | | | | | | |
|---|---|---|---|---|---|---|---|---|---|---|
| ***Allotype*** | ***1*** | ***2*** | ***3*** | ***4*** | ***5*** | ***6*** | ***7*** | ***8*** | ***9*** | ***10*** |
| ***1*** | 0 | | | | | | | | | |
| ***2*** | 0 | 19 | | | | | | | | |
| ***3*** | 0 | 2.6 | 0.2 | | | | | | | |
| ***4*** | 0 | 2.2 | 0 | 0 | | | | | | |
| ***5*** | 0 | 0 | 0 | 0 | 0 | | | | | |
| ***6*** | 0 | 0.4 | 0 | 0 | 0 | 0 | | | | |
| ***7*** | 0 | 13.4 | 1.2 | 0.8 | 0 | 0.4 | 3.6 | | | |
| ***8*** | 0 | 23.4 | 1.2 | 1.2 | 0 | 0.6 | 10 | 8.2 | | |
| ***9*** | 0 | 0 | 0 | 0 | 0 | 0 | 0 | 0 | 0 | |
| ***10*** | 0 | 5.4 | 0 | 0.6 | 0 | 0 | 1.2 | 4.4 | 0 | 0 |

| % in population (SAS) | | | | | | | | | | |
|---|---|---|---|---|---|---|---|---|---|---|
| ***Allotype*** | ***1*** | ***2*** | ***3*** | ***4*** | ***5*** | ***6*** | ***7*** | ***8*** | ***9*** | ***10*** |
| ***1*** | 0.4 | | | | | | | | | |
| ***2*** | 3.5 | 4.9 | | | | | | | | |
| ***3*** | 2.2 | 3.7 | 2.2 | | | | | | | |
| ***4*** | 0.8 | 0.4 | 0.6 | 0.0 | | | | | | |
| ***5*** | 1.2 | 3.5 | 3.3 | 0.8 | 1.6 | | | | | |
| ***6*** | 1.0 | 2.5 | 1.6 | 0.2 | 1.6 | 0.4 | | | | |
| ***7*** | 0.8 | 1.8 | 1.8 | 0.0 | 1.4 | 0.4 | 0.0 | | | |
| ***8*** | 3.5 | 9.8 | 8.4 | 1.0 | 5.1 | 3.1 | 2.7 | 8.4 | | |
| ***9*** | 0.0 | 0.0 | 0.0 | 0.0 | 0.0 | 0.0 | 0.0 | 0.0 | 0.0 | |
| ***10*** | 0.8 | 1.6 | 2.5 | 0.8 | 1.6 | 0.6 | 0.6 | 2.9 | 0.0 | 0.8 |

**Supplemental Table 6:** Frequency of ERAP1 1-10 allotype combinations in genetic samples that carry the [A,A],[A,B] and [B,B] ERAP1 allotypes. Numbers indicate % values. Cells are color coded (red=high, yellow=medium, green=low).

[A,A] ERAP2 (495 samples)

| ERAP1 allotype | 1 | 2 | 3 | 4 | 5 | 6 | 7 | 8 | 9 | 10 |
|---|---|---|---|---|---|---|---|---|---|---|
| **10** | 1.6 | 2.0 | 1.8 | 0.0 | 0.2 | 1.8 | 1.4 | 8.7 | 0.0 | 2.2 |
| **9** | 0.0 | 0.0 | 0.0 | 0.0 | 0.0 | 0.0 | 0.0 | 0.0 | 0.0 | 0.0 |
| **8** | 3.2 | 8.5 | 9.1 | 1.6 | 2.4 | 3.0 | 9.1 | 19.2 | 0.0 | 8.7 |
| **7** | 1.0 | 1.8 | 2.2 | 0.6 | 0.6 | 0.4 | 1.8 | 9.1 | 0.0 | 1.4 |
| **6** | 0.4 | 0.8 | 0.4 | 0.6 | 0.2 | 0.2 | 0.4 | 3.0 | 0.0 | 1.8 |
| **5** | 0.4 | 0.8 | 0.4 | 0.4 | 0.4 | 0.2 | 0.6 | 2.4 | 0.0 | 0.2 |
| **4** | 0.0 | 1.4 | 0.6 | 0.0 | 0.4 | 0.6 | 0.6 | 1.6 | 0.0 | 0.0 |
| **3** | 1.2 | 1.6 | 2.2 | 0.6 | 0.4 | 0.4 | 2.2 | 9.1 | 0.0 | 1.8 |
| **2** | 1.0 | 1.6 | 1.6 | 1.4 | 0.8 | 0.8 | 1.8 | 8.5 | 0.0 | 2.0 |
| **1** | 0.6 | 1.0 | 1.2 | 0.0 | 0.4 | 0.4 | 1.0 | 3.2 | 0.0 | 1.6 |

[A,B] ERAP2 (1086 samples)

| ERAP1 allotype | 1 | 2 | 3 | 4 | 5 | 6 | 7 | 8 | 9 | 10 |
|---|---|---|---|---|---|---|---|---|---|---|
| **10** | 2.0 | 5.2 | 2.8 | 0.6 | 1.7 | 2.1 | 1.6 | 4.9 | 0.0 | 1.4 |
| **9** | 0.1 | 0.2 | 0.1 | 0.0 | 0.0 | 0.1 | 0.0 | 0.1 | 0.0 | 0.0 |
| **8** | 3.1 | 17.2 | 3.1 | 1.6 | 4.1 | 2.8 | 3.4 | 3.8 | 0.1 | 4.9 |
| **7** | 1.1 | 5.4 | 0.9 | 0.6 | 0.7 | 1.0 | 1.6 | 3.4 | 0.0 | 1.6 |
| **6** | 0.4 | 1.9 | 1.7 | 0.2 | 1.0 | 0.3 | 1.0 | 2.8 | 0.1 | 2.1 |
| **5** | 0.6 | 2.9 | 1.7 | 0.5 | 0.4 | 1.0 | 0.7 | 4.1 | 0.0 | 1.7 |
| **4** | 0.3 | 1.1 | 0.9 | 0.0 | 0.5 | 0.2 | 0.6 | 1.6 | 0.0 | 0.6 |
| **3** | 1.4 | 3.7 | 0.7 | 0.9 | 1.7 | 1.7 | 0.9 | 3.1 | 0.1 | 2.8 |
| **2** | 1.4 | 4.9 | 3.7 | 1.1 | 2.9 | 1.9 | 5.4 | 17.2 | 0.2 | 5.2 |
| **1** | 0.9 | 1.4 | 1.4 | 0.3 | 0.6 | 0.4 | 1.1 | 3.1 | 0.1 | 2.0 |

[B,B] ERAP2 (650 samples)

| ERAP1 allotype | 1 | 2 | 3 | 4 | 5 | 6 | 7 | 8 | 9 | 10 |
|---|---|---|---|---|---|---|---|---|---|---|
| **10** | 2.0 | 5.4 | 0.8 | 1.4 | 2.0 | 1.4 | 0.2 | 1.1 | 0.0 | 0.8 |
| **9** | 0.0 | 0.0 | 0.2 | 0.0 | 0.0 | 0.0 | 0.0 | 0.0 | 0.0 | 0.0 |
| **8** | 1.7 | 7.1 | 0.3 | 0.3 | 1.4 | 0.8 | 0.2 | 1.4 | 0.0 | 1.1 |
| **7** | 0.0 | 5.4 | 0.6 | 0.8 | 0.5 | 0.2 | 0.3 | 0.2 | 0.0 | 0.2 |
| **6** | 2.8 | 5.5 | 0.3 | 0.8 | 1.7 | 1.1 | 0.2 | 0.8 | 0.0 | 1.4 |
| **5** | 1.2 | 3.7 | 2.0 | 0.6 | 1.4 | 1.7 | 0.5 | 1.4 | 0.0 | 2.0 |
| **4** | 1.5 | 3.7 | 0.0 | 0.3 | 0.6 | 0.8 | 0.8 | 0.3 | 0.0 | 1.4 |
| **3** | 0.9 | 6.0 | 1.4 | 0.0 | 2.0 | 0.3 | 0.6 | 0.3 | 0.2 | 0.8 |
| **2** | 5.5 | 22.3 | 6.0 | 3.7 | 3.7 | 5.5 | 5.4 | 7.1 | 0.0 | 5.4 |
| **1** | 1.4 | 5.5 | 0.9 | 1.5 | 1.2 | 2.8 | 0.0 | 1.7 | 0.0 | 2.0 |

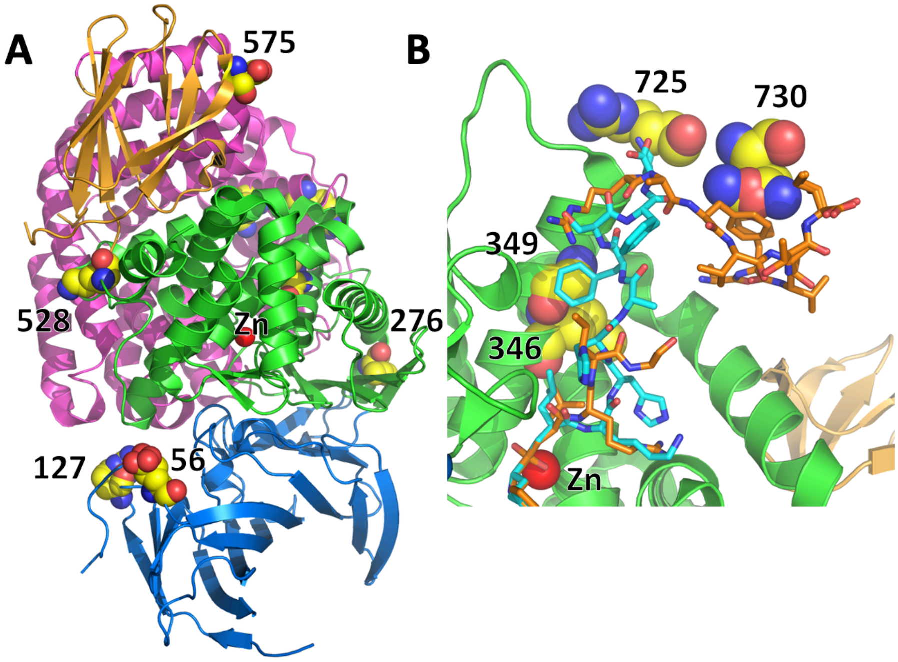


**Supplemental Figure 1:** Schematic representations of ERAP1 crystal structure (PDB codes 6RYF and 6RQX) indicating the positions of 9 SNPs. Structure is colored by domain (domain I in blue, domain II in green, domain III in orange and domain IV in magenta). Active site Zn(II) atom is shown as a red sphere). Panel A, polymorphic residues that lie on the outside of the protein are indicated by spheres (carbon=yellow, oxygen=red, nitrogen=blue). Panel B, polymorphic residues that lie in the inside of the substrate binding cavity of the enzyme are shown as spheres (carbon=yellow, oxygen=red, nitrogen=blue). Peptide substrates co-crystallized with ERAP1 are shown in stick representation (blue=10mer, orange=15mer).

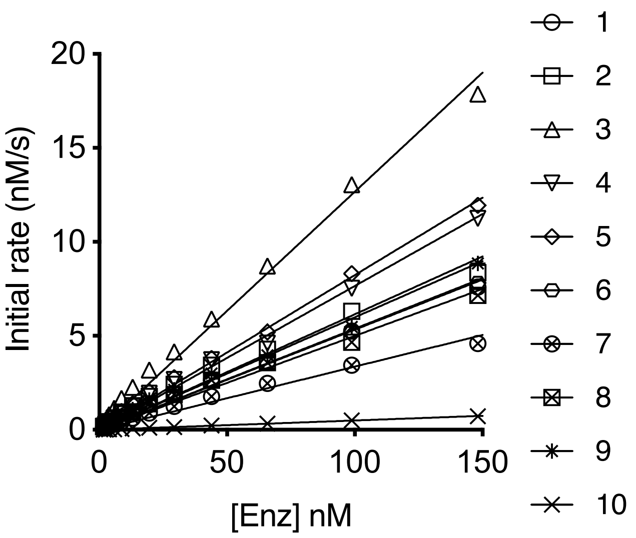


**Supplemental Figure 2:** Initial rates of hydrolysis of Leu-AMC by ERAP1 allotypes plotted against enzyme concentration.

**Supplemental Table 7:** Enzymatic parameters of each ERAP1 allotype versus small dipeptide substrates.

| Leu-AMC substrate | | | | Leu-pNA substrate | | |
|---|---|---|---|---|---|---|
| **Allotype** | **Sp. Act. ($s^{-1}$)** | **$k_{cat}/K_M$ ($M^{-1}\,s^{-1}$)** | **$V_{max}$ ($s^{-1}$)** | **Hill coeff** | **$K_{half}$ (mM)** | **$K_{prime}$ (mM)** |
| **1** | 0.0505±0.0011 | 2638 ± 32 | 2.95 ± 0.1 | 1.58 ± 0.09 | 0.85 ± 0.05 | 0.77 ± 0.08 |
| **2** | 0.0558±0.0025 | 2529 ± 18 | 2.8 ± 0.48 | 1.35 ± 0.31 | 0.95 ± 0.31 | 0.94 ± 0.42 |
| **3** | 0.1202±0.0026 | 4296 ± 49 | 4.98 ± 0.16 | 1.49 ± 0.08 | 0.8 ± 0.05 | 0.72 ± 0.08 |
| **4** | 0.0736±0.0019 | 3130 ± 15 | 10.47 ± 0.56 | 1.53 ± 0.07 | 1.7 ± 0.13 | 2.26 ± 0.19 |
| **5** | 0.0795±0.0014 | 3519 ± 29 | 8.07 ± 0.3 | 1.51 ± 0.06 | 1.4 ± 0.08 | 1.65 ± 0.12 |
| **6** | 0.0513±0.001 | 2273 ± 12 | 9.94 ± 0.79 | 1.43 ± 0.09 | 1.94 ± 0.23 | 2.57 ± 0.31 |
| **7** | 0.0311±0.0013 | 1706 ± 33 | 6.13 ± 0.35 | 1.46 ± 0.06 | 2.12 ± 0.17 | 2.99 ± 0.24 |
| **8** | 0.0463±0.0016 | 2492 ± 17 | 9.45 ± 0.35 | 1.6 ± 0.05 | 1.87 ± 0.1 | 2.72 ± 0.15 |
| **9** | 0.0568±0.0017 | 2747 ± 18 | 9.33 ± 0.32 | 1.55 ± 0.05 | 1.81 ± 0.09 | 2.52 ± 0.13 |
| **10** | 0.0049±0.0001 | 240 ± 5 | nd | 1.53 ± 0.08 | nd | nd |

**Supplemental Table 8:** Calculated parameters from Michaelis-Menten analysis of trimming of the 9mer peptide YTAFTIPSI by ERAP1 allotypes.

| | 9mer peptide | | |
|---|---|---|---|
| **Allotype** | **$k_{cat}$ ($s^{-1}$)** | **$K_M$ (μM)** | **$k_{cat}/K_M$ ($M^{-1}$ $s^{-1}$)** |
| **1** | 0.527 ± 0.014 | 6.5 ± 0.6 | 80523 ± 7251 |
| **2** | 0.553 ± 0.016 | 4.9 ± 0.5 | 113718 ± 12448 |
| **3** | 0.761 ± 0.021 | 13 ± 1 | 58480 ± 4601 |
| **4** | 0.655 ± 0.022 | 22.2 ± 1.8 | 29463 ± 2537 |
| **5** | 0.4 ± 0.015 | 14 ± 1.3 | 28478 ± 2938 |
| **6** | 0.403 ± 0.01 | 16.8 ± 1.1 | 23915 ± 1648 |
| **7** | 0.389 ± 0.027 | 25 ± 3.6 | 15567 ± 2466 |
| **8** | 0.489 ± 0.013 | 16 ± 1.1 | 30614 ± 2207 |
| **9** | 0.589 ± 0.018 | 23.8 ± 1.6 | 24717 ± 1826 |
| **10** | 0.051 ± 0.002 | 23.8 ± 1.4 | 2134 ± 144 |

**Supplemental Figure 3:** Bubble chart showing the frequency of allotype combinations in the five human populations analyzed color-coded by their estimated activity. Allotypes have been clustered based on activity (highest activity top right of each panel, lowest bottom left).

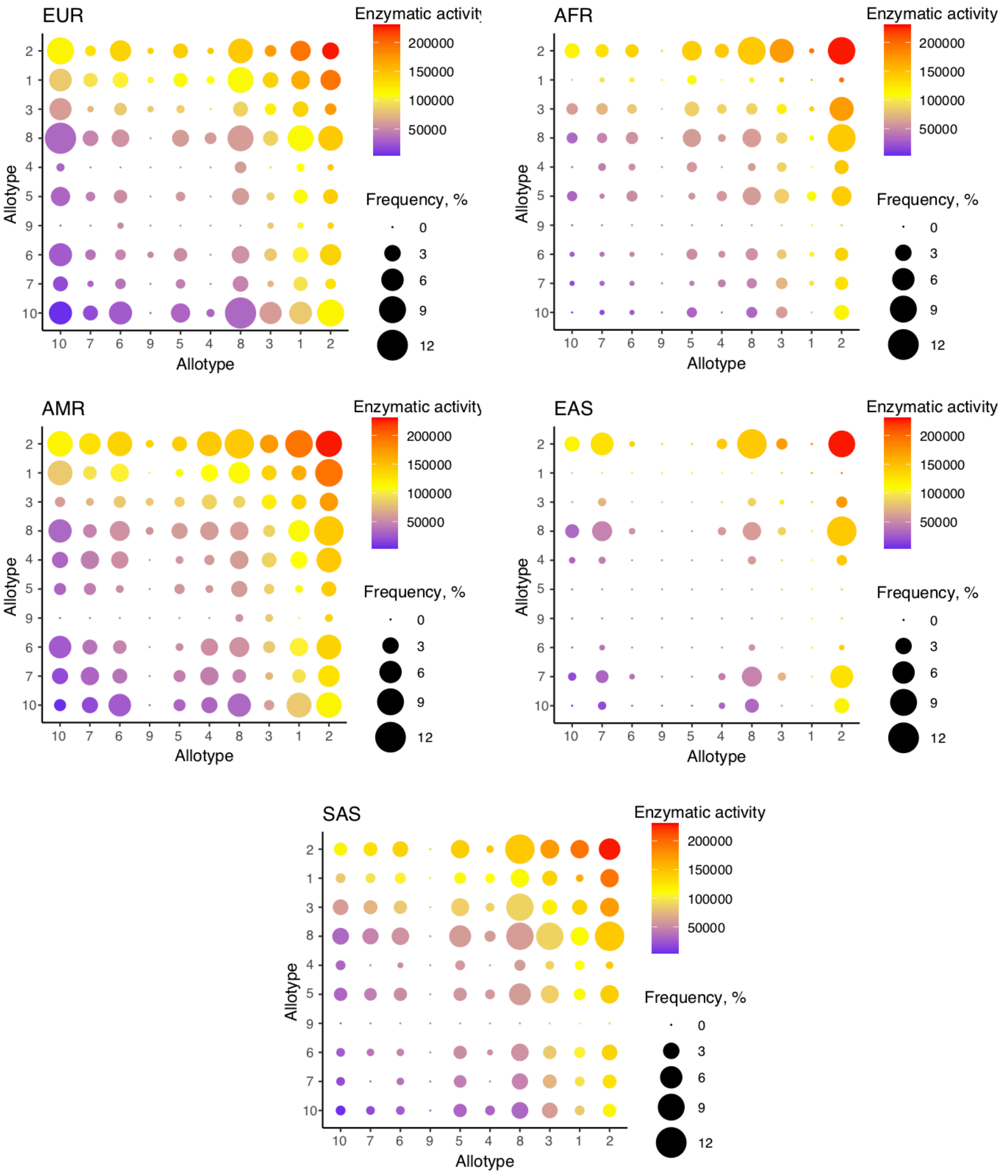
